# Enhancing rigor and reproducibility in maternal immune activation models: practical considerations and predicting resilience and susceptibility using baseline immune responsiveness before pregnancy

**DOI:** 10.1101/699983

**Authors:** Myka L. Estes, Kathleen Farrelly, Scott Cameron, John Paul Aboubechara, Lori Haapanen, Joseph D. Schauer, Aurora Horta, Kathryn Prendergast, Jeremy A. MacMahon, Christine I. Shaffer, Catherine T. Le, Greg N. Kincheloe, Danielle John Tan, Deborah van der List, Melissa D. Bauman, Cameron S. Carter, Judy Van de Water, A. Kimberley McAllister

**Affiliations:** Center for Neuroscience, University of California, Davis; Department of Internal Medicine, University of California, Davis; Department of Dermatology, University of California, Davis; Dept. of Psychiatry, University of California, Davis; Imaging Research Center, University of California, Davis

**Author notes:** Corresponding author: A. Kimberley McAllister, Ph.D. Professor & Director, Center for Neuroscience UC Davis, 1544 Newton Court, Davis CA 95618. **Author contributions**: MLE, KF, SC, JPA, LH, JDS, CIS, CTL, DJT, & DVL performed the research; MLE, SC, DJT, DVL, JVW, MDB, & AKM designed the experiments; MLE, KF, SC, JPA, AH, KP, JAM, GNK, DJT, & DVL analyzed the data. MLE, SC, JPA, & AKM wrote the manuscript. MDB, CSC, and AKM designed and wrote the grants that funded the research.

## Abstract

Despite the potential of rodent models of maternal immune activation (MIA) to identify new biomarkers and therapeutic interventions for a range of psychiatric disorders, their value is currently limited by issues of scientific rigor and reproducibility. Here, we report three sources of variability—the immunogenicity of the poly(I:C), the baseline immune responsiveness (BIR) of the females prior to pregnancy, and differences in immune responses in C57/B6 dams across vendors. Similar to the variable effects of human maternal infection, MIA in mice does not cause disease-related phenotypes in all offspring and the magnitude and type of maternal response, determined by a combination of poly(I:C) dose and BIR, predicts offspring outcome. Together, our results provide recommendations for optimization of MIA protocols to enhance rigor and reproducibility and reveal new factors that drive susceptibility of some pregnancies and resilience of others to MIA-induced abnormalities in offspring.

## Introduction

Epidemiological evidence links maternal infection to several neuropsychiatric disorders including autism (ASD) and schizophrenia (SZ) (1). Because mouse models of maternal immune activation (MIA) show substantial face and predictive validity for ASD and SZ (2, 3), they are invaluable for identifying new biomarkers and therapeutic interventions that could be effective during pre-symptomatic periods. However, despite the use of MIA mouse models in hundreds of laboratories world-wide, the value of these models is currently limited by major issues of reproducibility stemming from heterogeneous and sometimes opposing findings in MIA offspring (2, 4, 5). For an environmental model such as MIA to have construct validity for ASD and SZ, a quantifiable threshold of MIA that causes disease-relevant phenotypes must be established like the threshold for maternal infection in humans linked to ASD and SZ (4, 6, 7)(8). The poly(I:C) model was originally designed to cause elevations in maternal serum interleukin-6 (IL-6) comparable to those induced by influenza (9, 10). Elevation in maternal IL-6 is necessary and sufficient for inducing a range of disease-relevant phenotypes in MIA offspring (9). However, many laboratories using this model do not measure maternal serum IL-6, and those that do report widely varying levels that have been steadily decreasing over time even when using the same dose, timing, delivery route, and source of poly(I:C) (11). For example, initial studies that established the mouse MIA model using mid-gestational intraperitoneal injection of poly(I:C) reported 10,000-20,000pg/ml IL-6 in maternal serum 3hr post-injection, while a recent report found only a 350pg/ml increase using an identical MIA induction protocol (9, 12). Here, we identify several novel sources of variability in generating MIA models and propose a new approach to reduce heterogeneity and improve reproducibility in these models.

## Materials and Methods

### Animal Care and Use

All studies were conducted with approved protocols from the University of California Davis Animal Care and Use Committee, in compliance with NIH guidelines. Virgin C57BL/6N mice were purchased from Charles River (CR; Kingston, NY), Taconic (TAC; Hudson, NY) and C57BL/6J mice from Jackson (JAX; Sacramento, CA). Mice for MIA experiments were bred in house and maintained on a 12:12h light:dark cycle at 21 ± 1°C with food and water *ad libitum*. All mice were housed in Techniplast Sealsafe individually ventilated cages (IVC) with corncob bedding. Vivarium-specific environmental factors such as cage system (13), bedding (14), temperature and humidity (15), technician sex (16), and microbiota (17) exposure may all contribute to MIA susceptibility and outcomes (11) (**Supplemental Note 1**).

### Baseline immune responsiveness measurement

At 7 weeks of age, virgin female mice were injected intraperitoneally (IP) with a low dose of high molecular weight (HMW) Poly(I:C) dsRNA (InvivoGen, San Diego, CA. Cat# tlrl-pic). At 2.5hrs or 4hrs post-injection, whole blood from individual animals was taken via tail snip and spun at 14,000 rpm at 4°C for 8min. Serum was isolated and IL-6 levels were measured using a Bio-Rad Luminex system (Bio-Rad, Laboratories, Hercules, CA) according to the manufacturer’s protocol. The baseline immune responsiveness (BIR) of the dams was assessed by their relative serum interleukin-6 (IL-6) levels, and those levels were used to divide the dams into 3 groups (low, medium, and high) following priming and prior to breeding **(Supplementary Note 2)**. All injections were performed by female handlers.

### Maternal Immune Activation

As previously described (18), pregnant mice at gestational day GD12.5 were injected intraperitoneally (IP) with poly(I:C) dsRNA from Sigma Aldrich (St. Louis, MO, cat# P9582, #P1530, #P0913) and InvivoGen (cat# tlrl-picw and # tlrl-pic) at various doses spanning 20-200 mg/kg or with vehicle control (sterile 0.9% saline) injected at 5ul/ gram of body weight. Pregnancy was determined to be GD12.5 by visualizing vaginal plug (GD0.5) and by an increase in body weight. At time of injection, the male was separated from the dam. Serum was isolated from blood via tail snip for IL-6 levels at either 2.5 or 4hr post injection and serum was isolated from trunk blood 48hr after injection for IL-17a protein levels. Temperature, weight and sickness behavior were observed at injection, bleed and 24hr post injection times. Sickness behavior was measured as an index of maternal inflammation and scored using a subjective scale of increasing activity (from 1 to 3) of little to no movement or response to being handled to normal resistance to capture and restraint of the dam 2.5hr post injection. Experiments were performed on female mice before pregnancy, in pregnant dams, or in newborn or young adult offspring as indicated for each assay.

### Immunoblotting

Western blot analysis was used to investigate protein levels of STAT3, MEF2A, and tyrosine hydroxylase (TH) from the total homogenate of striatal (Str) brain samples. Striata from P0 male MIA offspring from the 3 BIR groups and controls were dissected in HBSS, frozen in liquid nitrogen, and stored at −80°C. Samples were disrupted using a probe sonicator (Qsonica Sonicator Q500) with an amplitude of 20% for 5sec in 2X Laemmli buffer, then denatured at 85°C for 5min. Lysates were centrifuged at 16,000g for 10min at room temperature. The supernatant was collected and stored at −80°C. Total protein content was measured using the Pierce BCA Protein Assay Kit (Thermo Fisher), using bovine serum albumin as the calibration standard. Dithiothreitol (Sigma D9779- 10G) was then added as a reducing agent to the samples at a final concentration of 100mM, and heated at 85°C for 2min before loading onto a gel. Equal amounts of protein (5ug/lane) were run under reducing conditions on 7.5% TGX gels (Bio-Rad) and electrophoretically transferred onto PVDF membranes (Bio-Rad 162-0177). Membranes were blocked with Odyssey blocking buffer (TBS; Li-Cor) and incubated with: i) monoclonal anti-STAT3 (12640S, 1:1,000; Cell Signaling); ii) monoclonal anti-MEF2A (ab76063, 1:1,000; AbCam); and (iii) polyclonal anti-TH (P40101-150, 1:1,000; Pel-Freez). After washing 3 times with TBS + 0.05% Tween20, membranes were incubated for 45min with fluorescent-tagged secondary antibodies (925-32213 and 925-68072, 1:15,000; Li-Cor). After 4 additional washes in TBS/Tween20, bands were visualized at 680nm and 800nm using the Odyssey CLx imaging system (Li-Cor). Results were standardized using β-tubulin, detected using anti-β-tubulin (Millipore MAB3408; 1:2,000). Band intensities were measured using Image Studio software (Li-Cor).

### Flow cytometry

Splenocytes were isolated from MIA or control dams 48h after injection. For intracellular cytokine assessment, mononuclear cells (1×10^6^ cells/mL) were cultured for 6 h with or without phorbol 12-myristate 13-acetate (PMA, 50ng/mL; Sigma), ionomycin (50ng/mL; Sigma), and GolgiStop (BD) in T cell media: RPMI 1640 (Invitrogen) supplemented with 10% (v/v) heat-inactivated FBS (Hyclone), and 2mM glutamine. Intracellular cytokine staining was performed according to the manufacturer’s protocol (Cytofix/Cytoperm buffer set from BD with PE-Cy7-conjugated anti-CD4 (RM4-5), eFluor 450-conjugated anti-TCR-β (H57-597), PE-conjugated anti-IL-17a (eBio17B7), FITC-conjugated anti-IFN-γ (XMG1.2), and fixable viability dye eFluro 780). For transcription factor assessment, isolated mononuclear cells were stained according to the manufacturer’s protocol (Fix/perm buffer set from eBiosciences with PE-Cy7-conjugated anti-CD4 (RM4-5), eFluor 450-conjugated anti-TCR-β (H57-597), PerCP-Cy5.5-conjugated anti-T-bet (4B10), AF488-conjugated anti-Gata3 (TWAJ), PE-conjugated anti-Foxp3 (FJK-16s), APC-conjugated anti-RORγt (B2D)). All antibodies were purchased from eBiosciences except for anti-IFN-γ, which was purchased from Biolegend. LSRFortessa (BD) and FlowJo software (Tree Star) were used for flow cytometry and analysis. Live, CD4^+^ TCRβ^+^ cells were assessed for cytokine and transcription factor expression. FC was dissected at P0 from the MIA and control offspring and acutely dissociated in papain. Dissociated cells were stained for APC-conjugated anti-ACSA-2 (IH3-18A3), anti-O4 (O4), and anti-CD11b (M1/70.15.11.5) (Miltenyi Biotec), the anti-MHCI subtype H2-Kb-PE (AF6-88.5) (BD) and a viability marker (Far Red Dead Cell Stain from Invitrogen) and then fixed in 1% PFA. Samples were run on a FACSCalibur (BD) flow cytometer and analyzed by Flowjo (Treestar). Data were gated on live, ACSA^−^ O4^−^ CD11b^−^ cells and assessed for sMHCI by anti-H2-Kb-PE fluorescence intensity.

### Neuronal culture

Neurons from postnatal day 0-1 (P0-1) male mouse frontal cortex (FC) were dissociated with papain and plated at a density of 50K/cm^2^ on poly-L-lysine coated glass coverslips (as previously described (19)) and maintained in GS21 media (MTI-Global).

### Immunocytochemistry (ICC)

Cells from FC f newborn offcpsing were fixed at 8 days in vitro (DIV) in 4% paraformaldehyde, 4% sucrose in PBS at 4°C for 10min. Neurons were permeabilized with 0.25% Triton X-100 for 5min, blocked with 10% BSA, and then incubated with primary and secondary antibodies diluted in 3% BSA. Primary antibodies used were: PSD-95 (K28/43, 1:1000; NeuroMab) and VAMP2 (104-202, 1:1000, Synaptic Systems). Secondary antibodies used were AF-488 and Cy3 conjugated anti-rabbit and anti-mouse (1:400, Invitrogen).

### Microscopy and image analysis

Imaging of neurons was performed using a Nikon C2+ (software v4.5) laser scanning confocal microscope with a 60X PlanApo oil immersion objective (1.4 NA). Laser power and PMT levels were held constant between groups within each experiment. 16-bit grayscale images for each channel were acquired sequentially and kalman-averaged over two scans at 2.5X artificial zoom. Synapses were defined as sites of at least 1 pixel of overlap between PSD-95 and VAMP2 puncta, identified using custom software (20). All images were prepared in Adobe Photoshop and Illustrator.

### Behavioral testing

P60-80 male offspring from the poly(I:C)-injected mothers (MIA offspring) were compared to offspring from saline-injected dams (controls). Mice were handled for three days prior to behavioral testing by male and female handlers. For handling, mice were acclimated in the testing room for 1hr prior to being gently handled for 3min followed by return to the home cage. For the self-grooming assay, the testing room was dimly lit (15-20 lux) and mice were acclimated for 1h. Each mouse was placed individually into an empty standard mouse cage (36×15×13cm) for 20min and behavior was video recorded. Videos were scored offline. After a 10min habituation period in the test cage, the behavior of each mouse was scored with a stopwatch for 10min for cumulative time spent grooming all body regions and for frequency of rearing events.

### qPCR for Segmented Filamentous Bacteria (SFB) detection

Bacterial genomic DNA was extracted from frozen fecal samples (within an experiment the samples were treated identically) following manufacturer’s directions using QIAamp Fast DNA Stool Mini kit (Qiagen, Hilden, Germany). Samples were quantified with a Real-Time PCR System (CFX connect Real-Time PCR, Bio-Rad) using fluorescent SYBR Green technology (Bio-Rad). PCR was performed with primers targeting 16s rRNA genes of SFB and eubacteria (EUB), results were quantified using the standard curve method. Levels of SBF DNA were normalized to EUB DNA and results presented as relative fold change compared to Taconic animals. The primer sequences for detection of SFB and EUB are as follows: SFB 5’- GACGCTGAGGCATGAGAGCAT -3’ and 5’- GACGGCACGGATTGTTATTCA- 3’; EUB 5’- ACTCCTACGGGAGGCAGCAGT- 3’ and 5’- ATTACCGCGGCTGCTGGC -3’.

### Data Analysis

For all experiments, data was collected from at least two (usually three) separate cohorts of animals and is presented as the mean ± SEM. Results were analyzed (Graphpad Prism v.7) by Student’s *t*-test, one-way ANOVA or two-way ANOVA, followed where appropriate by Tukey’s honestly significant difference *post hoc* test. For all graphs, significance was defined as: *p < 0.05, **p < 0.01, ***p < 0.001, ****p < 0.0001. When normalized, data were plotted relative to the appropriate, prep-matched control. Data were plotted in Graphpad Prism (v.7) and exported to Adobe Illustrator for figure preparation.

## Results

### Variability in the immunogenicity of different forms and lots of poly(I:C)

Recently, we discovered that the potency of most of the available sources of the potassium salt poly(I:C) used in the original MIA models had dramatically decreased, eliciting an IL-6 response in pregnant mice ranging from 0-2% of the originally reported MIA values (**Supplemental Figure 1**). This reduced potency appears to be caused by variability in the poly(I:C) itself because there is wide variation in dsRNA concentration and molecular weight (MW) across manufacturers, lots, and even bottles from the same lot (7)(4). This variability contributes to heterogeneous maternal immune responses (7) since poly(I:C) initiates viral responses in a dsRNA length-dependent manner (9, 21, 22). Pure poly(I:C) showed the least variability between lots in these measures and thus, is likely to result in less immunogen-induced variability in the model.

In order to identify the effective dose of our specific lot of pure poly(I:C), multiple doses were tested in pregnant C57/B6 mice (Charles River; CR). Enhanced sickness behavior was observed in all pregnant mice injected with 20, 30, and 40mg/kg of poly(I:C) at 4hr post-injection (**Figure 1a**) but, in contrast to previous reports, dams showed a dose-dependent *decrease* in temperature at this time-point, with only the 30mg/kg group reaching significance (**Figure 1b**). Importantly, no dose of poly(I:C) elicited an average IL-6 post-injection response of 10,000pg/ml at 4hr post-injection as reported in the initial MIA studies (9). However, when sampling at the apex of the IL-6 response (**Supplemental Figure 2**), dams receiving 30 or 40 mg/kg of poly(I:C) met this threshold and lost weight in response to immune challenge (**Figure 1c-d**). Despite being statistically significant, dams treated with the standard dose (20 mg/kg) fell below this IL-6 threshold on average (**Figure 1c**), trended toward weight gain rather than loss (**Figure 1d**), and did not show changes in temperature (**Figure 1b**). Dose-dependent effects were also observed for litter resorptions (**Figure 1e**). As in our previous report (19), synapse density was significantly decreased in cultures from MIA offspring from dams treated with all doses of poly(I:C) (**Figure 1f-g**), but levels of major histocompatibility complex I (MHCI) levels were significantly increased only by a 30 mg/kg dose (**Figure 1h**). The 30 mg/kg dose was selected as the most effective dose of our particular lot of pure poly(I:C) because it was the lowest concentration that minimized litter loss while eliciting reproducible changes in the biological measures obtained when poly(I:C) was more immunogenic (19). Together, our results indicate that the effective dose of poly(I:C) will differ between lots and should be determined for each new lot when generating the MIA model.

**Figure 1.**
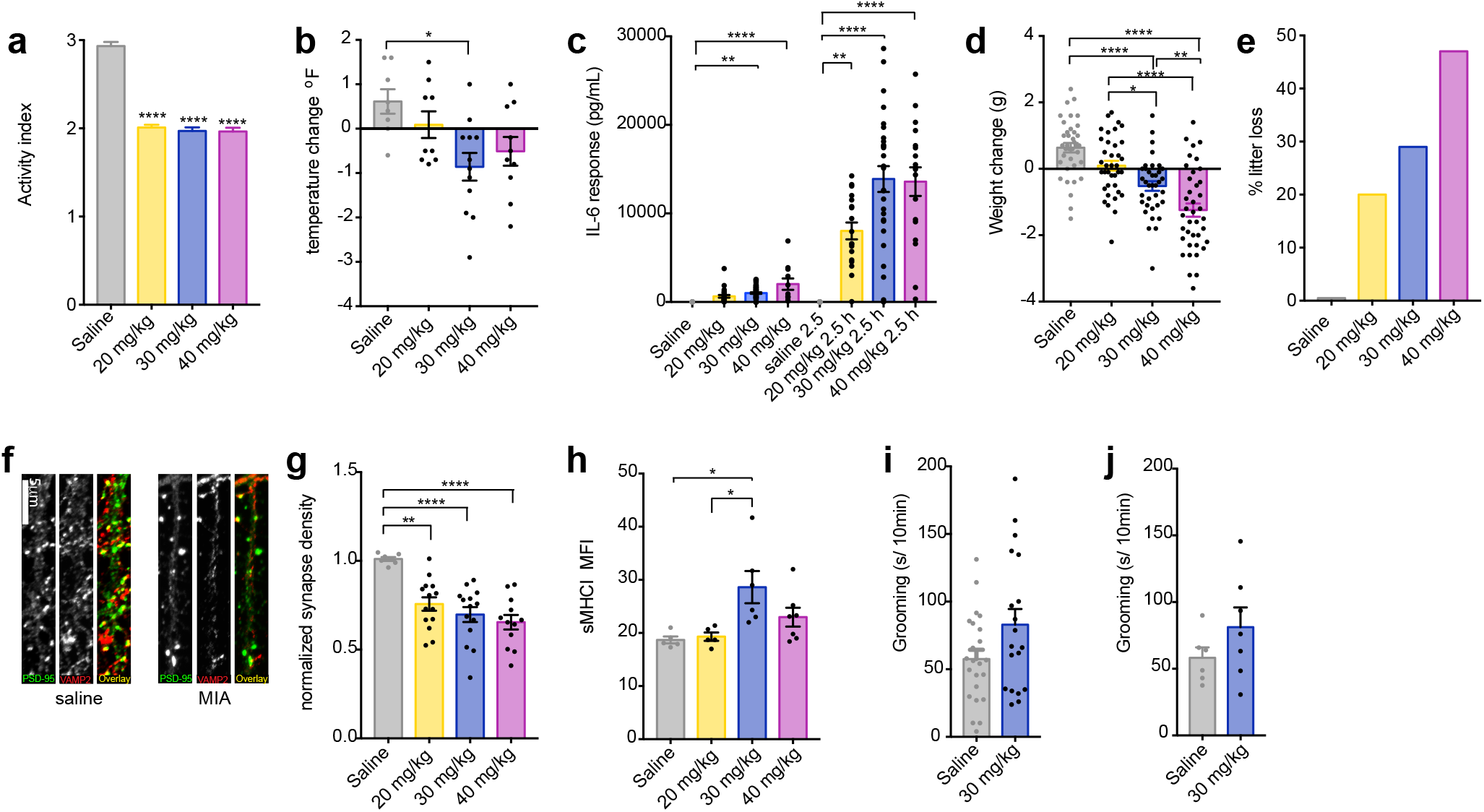
Measures of sickness behavior, litter loss, maternal IL-6, and offspring neuronal sMHCI are predictive of a disease-relevant dose of poly(I:C) (**a**) Sickness behavior in pregnant females at gestational day (GD) 12.5 was observed 4 hr post-injection with all 3 poly(I:C) doses—20, 30, and 40mg/kg (*F*_3,184_ = 102.3, *P* < 0.0001). (**b**) However, body temperature of GD12.5 dams 4hr post-injection was decreased at 30 and 40mg/kg, but not 20mg/kg, compared to saline, with only the 30mg/kg dose reaching significance (*F*_3,35_ = 4.289, *P* < 0.05). (**c**) Although all 3 doses caused significant elevations in maternal serum IL-6 at 2.5 and 4 hr post-injection, IL-6 levels were much higher at 2.5hr than at 4hr post-injection and only the 30 and 40mg/kg doses reached the 10,000 pg/ml IL-6 MIA threshold (*F*_3,35_ = 25.54, *P* < 0.0001). (**d**) The 30 and 40, but not 20, mg/kg doses caused significant weight loss in dams 24hr after poly(I:C) injection compared to saline (*F*_7,187_ = 26.93, *P* < 0.0001). (**e**) Poly(I:C) caused litter loss in a dose-dependent manner. (**f**) Representative images of glutamategic synapses on dendrites cultured from the frontal cortex (FC) of newborn offspring of saline-injected (saline) or poly(I:C)-injected (MIA) mothers. Neurons were immunostained at 8 DIV for excitatory synapse density using antibodies against PSD-95 (*green*) and VAMP2 (*red*). Scale bar = 5 μm. (**g**) All doses of poly(I:C) were associated with a significant decrease in synapse density (SD) (*F*_3,43_ = 11.01, *P* < 0.0001). Values were normalized to saline control (*n* ≥ 7 litters). (**h**) Surface MHCI (H2-Kb) was significantly increased on acutely dissociated neurons from FC of P0 offspring following MIA elicited by 30, but not 20 or 40, mg/kg poly(I:C), as assessed using flow cytometry (*n* ≥ 5 mice per group, 2 experiments) (*F*_3,19_ = 5.156, *P* < 0.01). (**i-j**) The effects of poly(I:C) dose on male offspring grooming (secs/10 min) behavior were assessed between P60-80. (**i**) Although there was a strong trend toward increased grooming in the MIA offspring from the 30 mg/kg group, the results were not significant due to the high variance (*P* = 0.06) (*n* ≥ 19 mice per group from at least 6 litters, 3 experiments). (**j**) The lack of significance and high variance remained after accounting for litter effects by averaging the grooming behavior of individuals within each litter (*P* = 0.22) (*n* ≥ 6 litters, 3 experiments). Bars represent mean ± s.e.m *p < 0.05, **p < 0.01, ***p < 0.001.

### Baseline immune response in dams prior to pregnancy predicts susceptibility or resilience of offspring to MIA-induced phenotypesg

MIA offspring display a range of aberrant behaviors with relevance for neurodevelopmental disorders, including perseverative behaviors (23–25). Although there was a trend towards increased grooming in MIA offspring using 30mg/kg pure poly(I:C), the difference was not statistically significant due to the high amount of variance (**Figure 1i**), which was also evident when we accounted for litter effects (**Figure 1j**) (26). Because the variance of the immune response of pregnant mice to poly(I:C) was also large (**Figure 1c**), we tested if individual differences in the baseline immune response (BIR) of the dams could account for the heterogeneity in MIA outcomes in offspring. Prior to mating, virgin females were injected with a low dose (4 mg/kg) of poly(I:C) and serum IL-6 levels were measured 2.5 hr post-injection (**Supplemental Note 2**). The mice were partitioned into three BIR groups based on the IL-6 responses across the cohort—low (first quartile), medium (second and third quartile), and high (fourth quartile) responders. Because individual IL-6 responses were similar in magnitude after a second poly(I:C) challenge, the BIR of females appears to be a stable trait (**Supplemental Fig. 3**; **Supplementary Note 2**). Importantly, grouping MIA offspring based upon BIR unmasked a significant increase in grooming in MIA offspring from low and medium BIR groups (**Figure 2a**). Unexpectedly, MIA offspring from high BIR dams showed no difference from controls. A different pattern across BIR groups was evident for abnormalities in rearing, an exploratory behavior in rodents indicative of anxiety (27, 28). Medium and high BIR MIA offspring showed a significant difference in rearing compared to control offspring (**Figure 2b**), but the direction of change was opposite in these groups—decreased in the medium and elevated in the high BIR MIA offspring (**Figure 2b**). There were no significant differences in either behavior in MIA offspring from any BIR group injected with 20 or 40 mg/kg poly(I:C), paralleling the dose-specific changes in maternal temperature and weight loss and offspring MHCI levels (**Supplemental Figure 4**). These data suggest that a threshold of MIA must be crossed for phenotypes in offspring to be robust and that the level of MIA above this threshold can lead to distinct combinations of aberrant phenotypes in offspring, reminiscent of the wide range of outcomes in humans exposed to maternal infection (2, 29).

**Figure 2.**
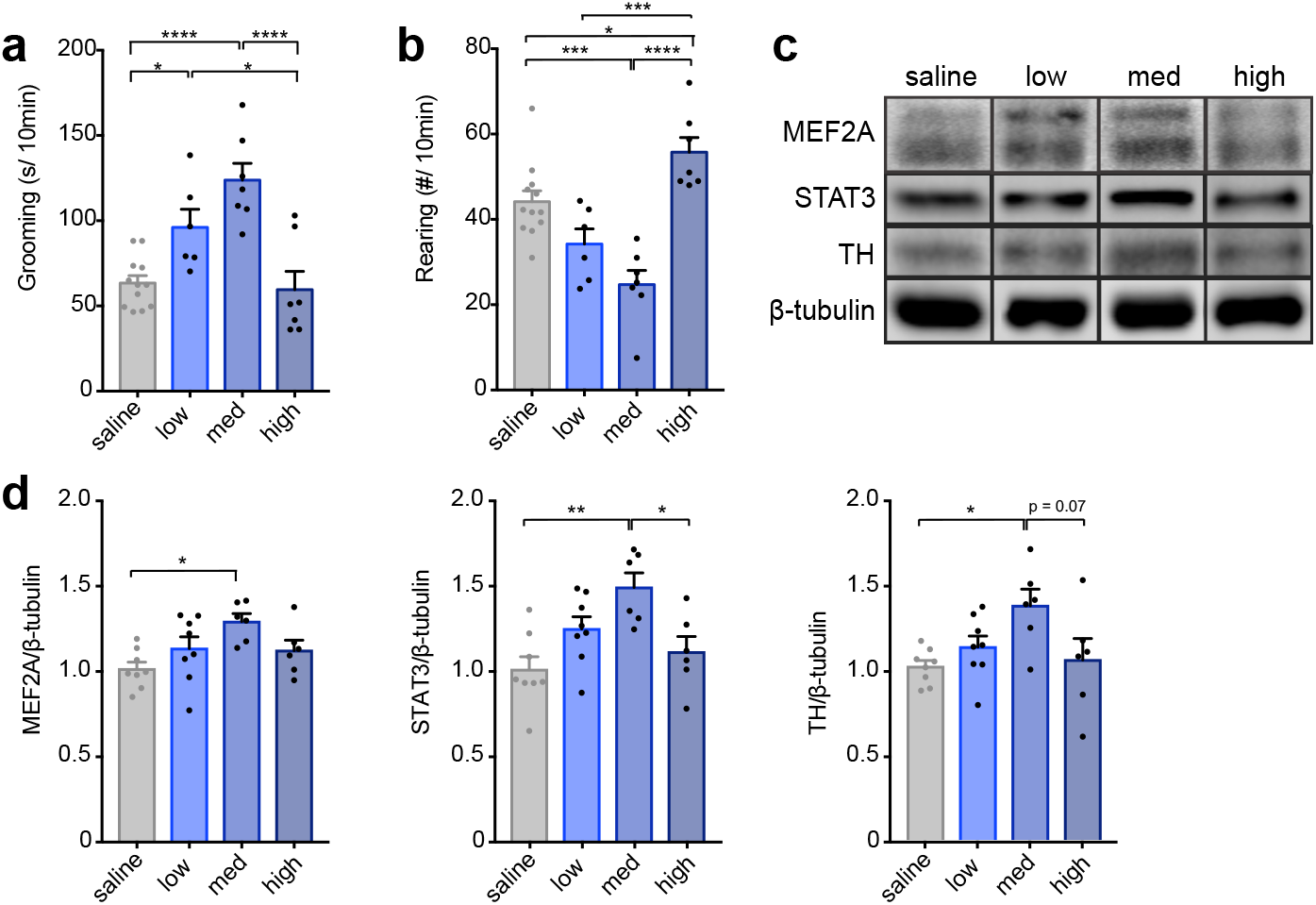
Baseline immune responsiveness of females prior to pregnancy determines MIA outcomes in offspring. Grouping litter by BIR of the dam revealed BIR-dependent changes in (**a**) grooming and (**b**) rearing behavior of young adult male offspring following MIA induced by 30mg/kg poly(I:C). (**a**) Time spent grooming was significantly increased in the low and medium BIR groups (*F*_3,28_ = 16.06, *P* < 0.0001), but there was no difference from controls in the high BIR group (*P* = 0.98). **(b**) For rearing, the medium BIR group showed a significant decrease in frequency, while the high BIR group showed the opposite behavior—a significant decrease compared to controls (*F*_3,28_ = 13.5, *P* < 0.0001). Low BIR MIA offspring were not different from controls. (**c**) Representative western blots showing increased MEF2, STAT3, and TH protein from striatum of MIA offspring compared to saline control offspring in a manner dependent on BIR of the dams. (**d**) Densitometry shows increased levels of MEF2A, STAT3, and TH protein relative to β-tubulin loading controls. (MEF2A: *F*_3,24_ = 3.968, *P* < 0.05; STAT3: *F*_3,24_ = 6.401, *P* < 0.01; TH: *F_3,24_ =* 3.668, P < 0.05). Data were averaged from two males per litter and 6-7 litters per BIR group. Bars represent mean ± s.e.m *p < 0.05, **p < 0.01, ***p < 0.001.

A significant advance for the field would be identification of biomarkers with high predictive value for specific MIA outcomes in offspring. We identified three potential biomarkers in the striatum, a brain region involved in regulating perseverative behaviors (30), that correlate with MIA outcomes in offspring. MEF2A—a transcription factor that mediates synaptic deficits caused by MIA (19), STAT3—a major signaling protein downstream of cytokines, and tyrosine hydroxylase (TH)—an enzyme involved in the synthesis of dopamine—were increased selectively in newborn offspring from medium BIR MIA dams (**Figure 2c-d**). Regardless of whether future experiments show that these proteins are causal for MIA-induced repetitive behaviors, levels of these proteins in the striatum of offspring can be used to determine the effective dose of a new lot of poly(I:C) when access to behavioral or synaptic assays is limited.

### MIA engages distinct molecular pathways in C57/B6 mice from different vendors

Surprisingly, the source of mice of the same strain also plays a major role in determining whether offspring will be affected. The MIA model was historically generated with C57/B6 mice sourced from Charles River (CR) or Jackson (JAX) (31). However, it has recently been shown that a maternal microbiome component (SFB), found in C57/B6 mice sourced from Taconic (TAC), but not JAX, is necessary for the generation of MIA outcomes (17). In these MIA TAC mice, increased maternal IL-6 acts to stimulate the secretion of IL-17a from maternal T helper (Th) 17 cells, which is necessary and sufficient to cause aberrant behaviors in offspring (12). While this study did not find MIA phenotypes in offspring from SFB-negative JAX dams, mice from JAX have been used successfully as MIA models in previous reports (6, 9, 23, 32, 33). In order to determine whether SFB and IL-17a are required in the generation of MIA outcomes, we first confirmed that C57/B6 mice purchased from TAC harbor SFB while those from JAX do not and we found that CR mice have levels of SFB comparable to TAC mice (**Supplemental Fig. 5**). While baseline IL-17a levels were not significantly different in saline-injected CR and TAC dams, only TAC dams showed an increase in IL-17a 48 hr after poly(I:C) injection (**Figure 3a**). In response to immune challenge, splenic maternal T_H_17 cells increased significantly in TAC, but not CR dams (**Figure 3b**). In contrast to CR dams, TAC dams displayed no fetal loss in response to poly(I:C) suggesting that the cytokine changes caused by MIA that contribute to fetal loss may be distinct between C57/B6 mice sourced from CR vs. TAC. Additionally, while CR dams lost weight 24hr following poly(I:C) injection, TAC dams generally showed no weight loss or trended towards weight gain (**Figure 3c**). These surprising differences in maternal response to immune challenge dictated by the source of the same strain of mice may also contribute to differences in MIA outcomes.

**Figure 3.**
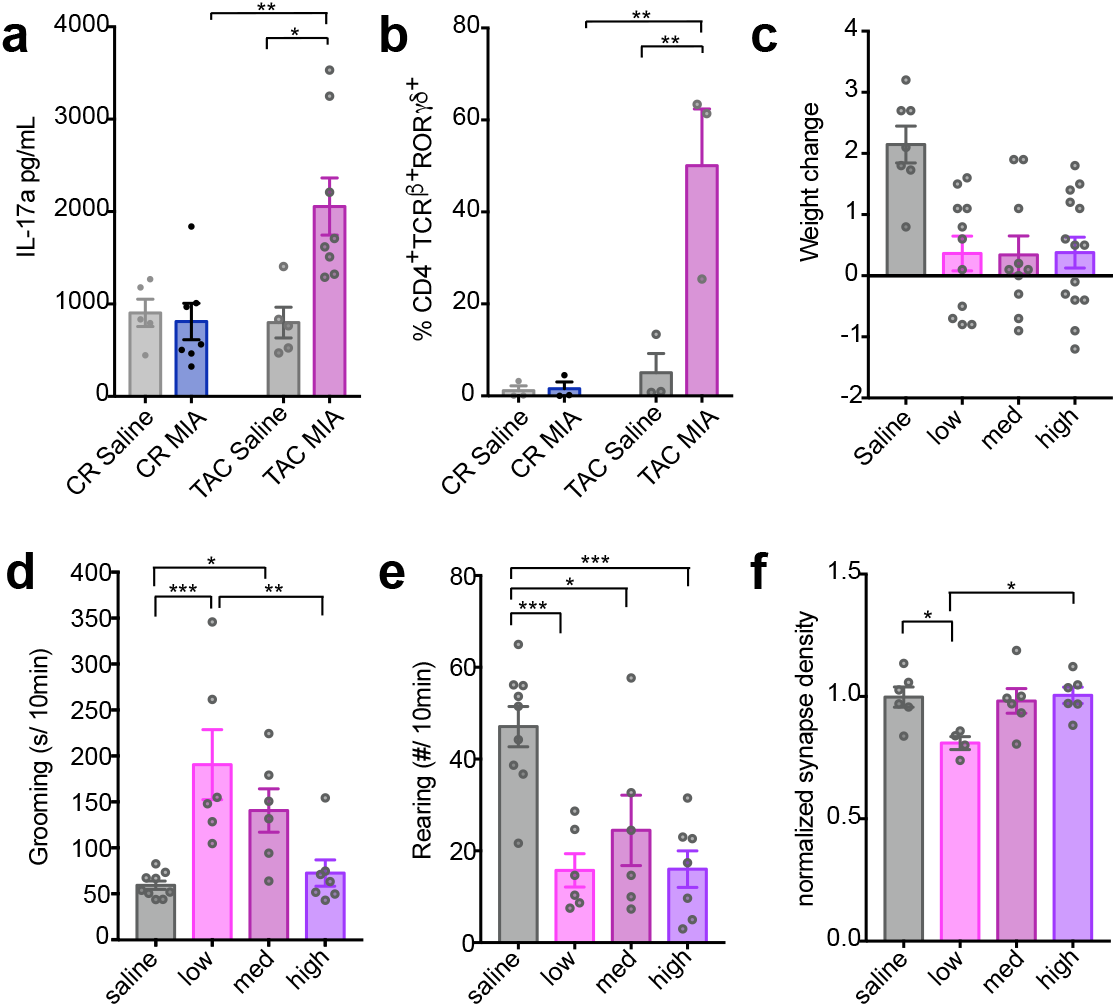
C57/B6 mice from different sources show altered susceptibilities to MIA. (**a**) Poly(I:C) injection at GD 12.5 caused dramatic elevations in serum concentrations of maternal IL-17a at GD14.5 in TAC, but not in CR, dams (significant interaction condition x source; *F*_1,21_ = 7.164, *P* < 0.05; *n* ≥ 5 mice per condition, > 2 experiments). (**b**) TAC but not CR dams show a significant increase in splenic T_H_17 cells 48hr following poly(I:C) injection (*F*_1,8_ = 11.48, *P* < 0.01). Splenic CD4^+^ TCRβ^+^ T cells stained intracellularly for RORγt from GD14.5 dams injected with poly(I:C) or saline at GD12.5 were detected using flow cytometry. (**c**) TAC dams do not lose weight in any BIR MIA group 24hr following poly(I:C) injection. MIA offspring from TAC dams treated with a 30 mg/kg dose were assessed for (**d**) grooming and (**e**) rearing behavior. MIA TAC offspring showed elevated grooming in low and medium BIR groups, but not the high BIR group (*F*_3,24_ = 8.82, *P* < 0.001). The effect size for time spent self-grooming in MIA offspring from low BIR dams was larger when animals were sourced from TAC (source x BIR: *F*_3,51_ = 3.81, *P* < 0.05; post hoc TAC low > CR low; *P* < 0.01). (**e**) Rearing was decreased in all BIR MIA groups in TAC mice (*F*_3,24_ = 9.984, *P* < 0.001). (**f**) Neurons from neonatal low BIR MIA offspring from TAC dams showed a significant decrease in synapse density (SD) (*F*_3,18_ = 4.014, *P* < 0.05). Values were normalized to saline control (*n* ≥ 4 litters). Bars represent mean ± s.e.m *p < 0.05, **p < 0.01, ***p < 0.001.

To test the effects of vendor on vulnerability of C57/B6 mice to MIA insult, we performed the same behavioral and synaptic assays on MIA and control offspring from dams sourced from TAC. Baseline grooming and rearing behavior was the same between control offspring from CR and TAC dams. Similar to MIA CR offspring, MIA TAC offspring from low and medium BIR dams showed increased self-grooming (**Figure 3d**). When comparing grooming behavior from MIA offspring we found a significant interaction between source and BIR and a larger effect size in time spent self-grooming in MIA offspring from low BIR dams when animals were sourced from TAC (source x BIR: *F*_3,51_ = 3.81, *P*<0.05; post hoc TAC low>CR low; *P*<0.01). Rearing behavior was also altered in TAC MIA offspring (**Figure 3e**). In contrast to CR MIA offspring, rearing was decreased in TAC MIA offspring from all BIR dams. There was a significant interaction between source and BIR (*F*_3,51_ = 6.445, *P*<0.001). Finally, there was also a significant decrease in the ability of neurons from MIA offspring of low BIR (but not medium or high) dams to form synapses (**Figure 3f**).

## Discussion

Issues of reproducibility of the MIA model are currently a significant obstacle for the field and hamper the use of this important translational model for pre-clinical research (2, 7, 11). Here, we identify several major sources of variability in the model: poly(I:C) immunogenicity and BIR of female mice within and between colonies of the same strain from different sources. Our data suggest that these factors contribute to discrepancies and must be considered when evaluating and comparing published results in the MIA field. Because poly(I:C) immunogenicity varies so much between forms and lots, we recommend that laboratories use the pure form of poly(I:C) to minimize within-lot variability and that they optimize the effective dose of poly(I:C) for each new lot in order to elicit a comparable level of MIA across lots and laboratories. Reporting measures of maternal cytokines are essential for any study using this model to allow for comparison across studies and maternal serum IL-6, while not the only important cytokine for MIA induction, is the cytokine that most labs use. Maternal levels of serum IL-6 of ~10,000pg/ml on average measured at 2.5hr post-injection appear to be a reliable threshold for induction of MIA leading to reproducible disease-like phenotypes in offspring. Because the optimization process is costly, labs might consider purchasing a large amount of a single lot of poly(I:C) to maximize the investment in establishing parameters for reproducible results. Finally, although it is well known that MIA phenotypes differ between mouse strains, our results indicate that the source of the mice is also a critical consideration for establishing the model since MIA activates distinct immune responses in C57/B6 mice from different vendors.

Even when controlling for the dose, delivery route, and MW of poly(I:C), MIA leads to a wide range of outcomes in offspring. Rather than being a problem for the model, this variability strengthens the construct and face validity of the model since one of the most important aspects of maternal viral infection in humans is the variability in outcomes and the fact that many pregnancies do not result in disease in offspring. Current approaches using MIA mouse models ignore this important consideration. For human disease, understanding the factors that drive susceptibility of some pregnancies and resilience of others to MIA-induced abnormalities in offspring is paramount.

Although variable outcomes in humans to disease risk factors are typically attributed to genetic differences, we found that BIR of female mice before pregnancy with the same genetic background may be a critical factor dictating susceptibility and resilience to MIA. Young C57/B6 virgin female mice exhibit a surprisingly wide range of BIR to a low-dose poly(I:C) challenge. Although a full characterization of the differences and causes underlying this range of BIR in cage-mates remains an important goal for the future, it is clear that BIR before pregnancy predicts MIA outcomes and determines the distinct constellation of phenotypes in offspring. Low BIR provides resilience to many of the effects of MIA, while medium BIR leads to greater perseverative behaviors and high BIR predicts more anxiety-like behaviors in offspring. Studying the circuit and molecular mechanisms underlying these distinct phenotypes caused by the same environmental insult could help to uncover factors that protect against, or increase susceptibility, to specific MIA outcomes. Our data also suggest that BIR should also be considered in human epidemiological study design in order to potentially unmask larger effect sizes and identify biomarkers for vulnerable populations.

---

**Supplemental Figure 1:**
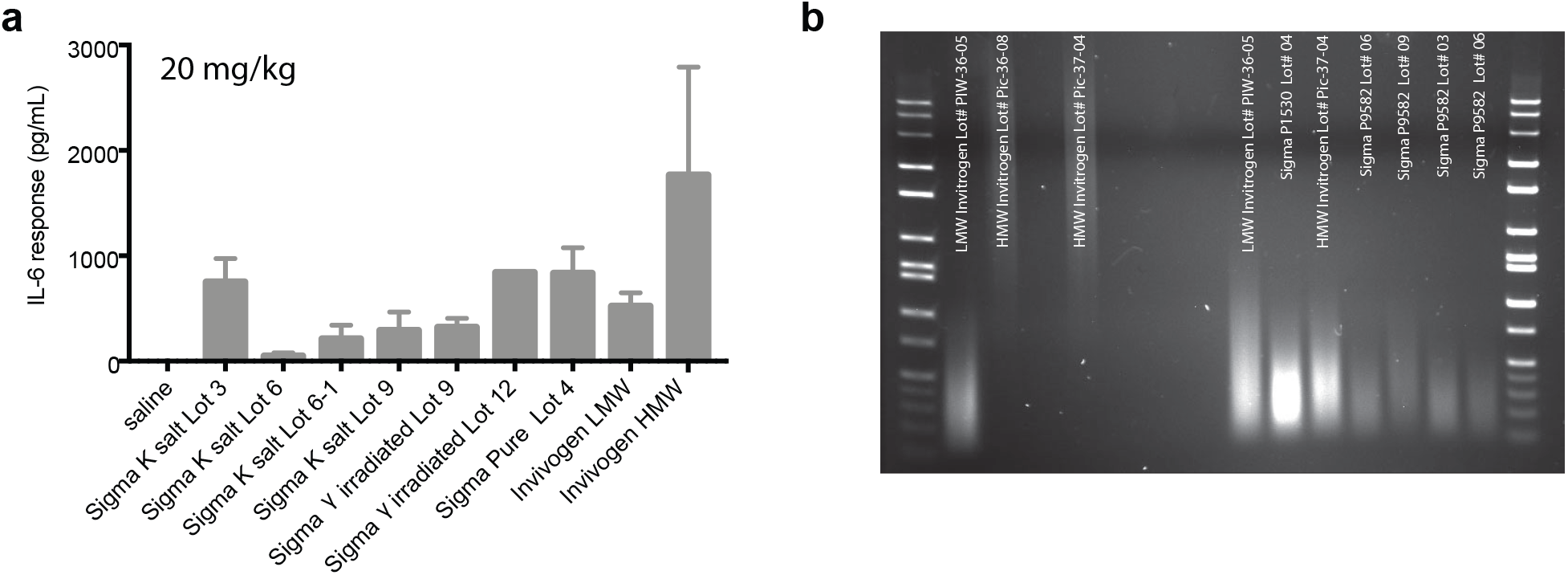
Variations in Poly(I:C) contribute to maternal IL-6 response. Lot number, bottle, and formula of poly(I:C) contribute to the measured IL-6 response. (**a**) Dams were injected with a 20 mg/kg dose of different forms of poly(I:C) as indicated at GD12.5 and trunk blood samples were taken 4hr post-injection. Serum IL-6 was measured using Luminex. All 4 lots tested (labeled 3, 6, 9, and 12) of the standard poly(I:C) used in the field, Sigma potassium salt poly(I:C) (cat# P9582), labeled Sigma K-salt, increased maternal IL-6 compared to saline although the magnitude of IL-6 elevation differed substantially between lots. There was also bottle-to-bottle variability in immunogenicity within a single lot (2 bottles of lot 6 were tested; labeled 6 and 6-1; bottle 6-1). Gamma-irradiation of the potassium salt did not alter immunogenicity within a lot as shown for lots 9 and 12 (lot 9 γirradiated; lot 12 γirradiated). Sigma pure poly(I:C) caused elevations in IL-6 to the maximal level as any of the potassium salt forms (lot 4). Poly(I:C) is also available from InvivoGen, which sells either low MW (LMW) or high MW (HMW) poly(I:C). While both forms elevated maternal IL-6 significantly compared to controls, dsRNAs of differing MW are known to initiate distinct immune responses and therefore may not be as similar to the general viral-like immune response initiated by the mixed MW Sigma compounds. Moreover, subsequent experiments using a second lot of HMW InvivoGen poly(I:C) caused complete litter loss in most mice injected, indicating either widely differing immunogenicity between lots or endotoxin contamination. (**b**) The concentration and molecular weight of dsRNA differs between lots and bottles of poly(I:C). 0.1 μg of poly(I:C) was run on a 1.5% TAE agarose gel. Each lot and even different bottles of the same lot differed in the molecular weight and concentration of dsRNA.

**Supplemental Figure 2:**
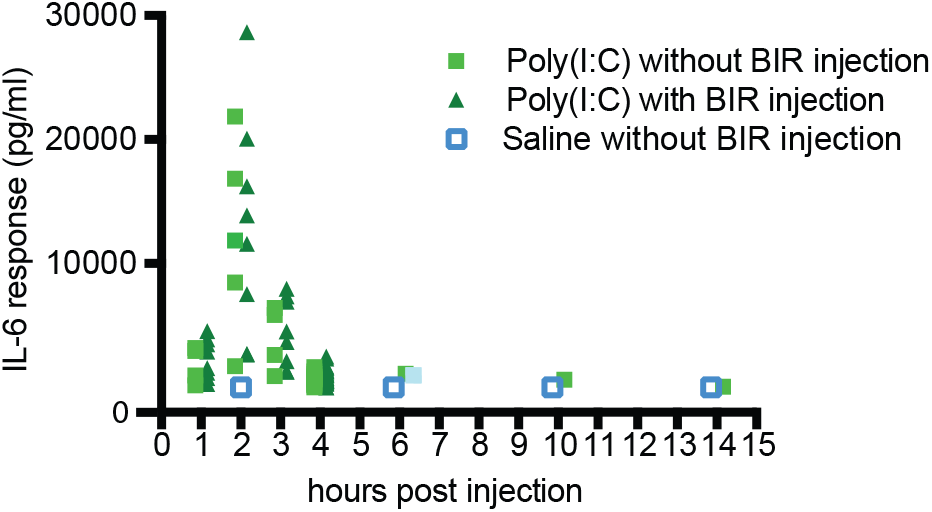
Time-course of maternal IL-6 response to 30 mg/kg poly(I:C) in dams that received a low dose poly(I:C) or no injection before mating. Dams were tail-bled at multiple time points following poly(I:C) or saline injection at GD 12.5 and isolated serum was analyzed by Luminex for IL-6. Maternal IL-6 peaked between 2 and 2.5hr post-injection for all dams regardless of whether they had received the low dose injection prior to mating. Pre-pregnancy low-dose poly(I:C) did not affect the kinetics of maternal IL-6 in response to poly(I:C). Saline injection at GD 12.5 did not alter maternal serum IL-6 at any time-point measured.

**Supplemental Fig. 3:**
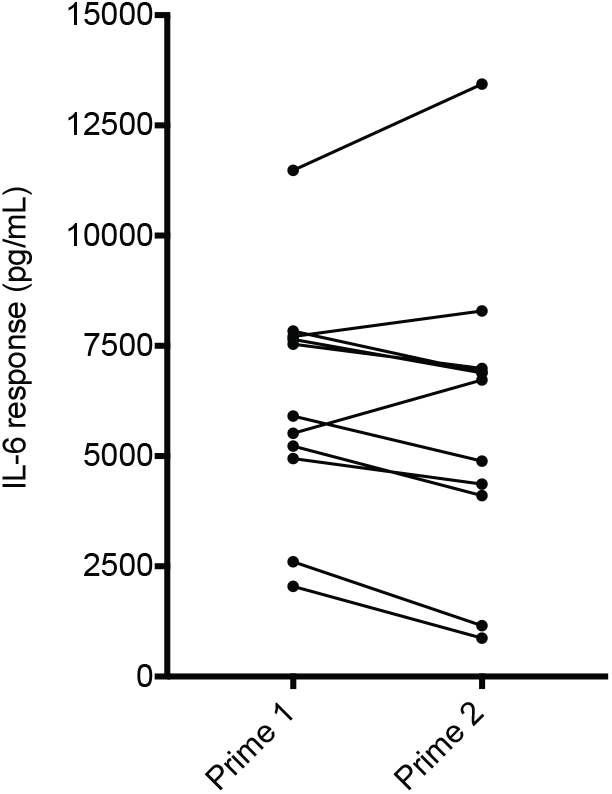
Baseline immune response to low dose Poly(I:C) is a stable trait in virgin females. 8 week-old virgin females were injected with low dose poly(I:C) and assessed for serum IL-6 by tail bleed 2.5hr post-injection. One week later, the same females were injected again with the same low dose of poly(I:C) and assessed for serum IL-6. All animals remained within their initial ‘low’, ‘medium,’ and ‘high’ BIR designation from the first to the second injection of low dose poly(I:C).

**Supplemental Fig. 4:**
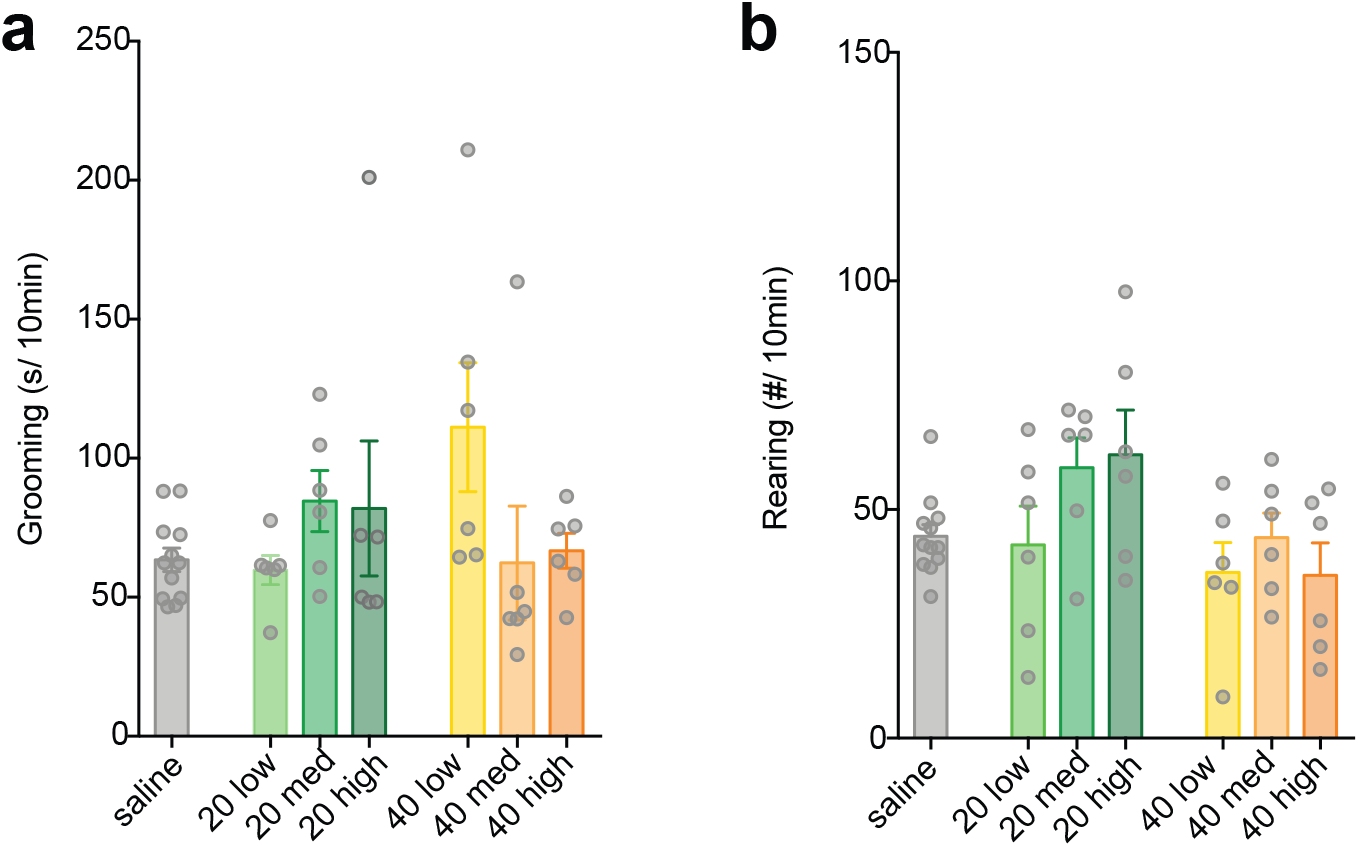
MIA offspring from dams treated with 20 and 40 mg/kg dose of poly(I:C) do not show MIA behavioral abnormalities, regardless of BIR. Not all doses of poly(I:C) produce MIA behaviors. No significant effect of 20 or 40 mg/kg poly(I:C) was observed on **a)** grooming (one-way ANOVA; 20 mg/kg: *F*_3,26_ = 1.153, *P* = 0.356; 40 mg/kg: *F*_3,26_ = 2.888, *P* = 0.055) or **b)** rearing (one-way ANOVA; 20 mg/kg: *F*_3,26_ = 2.561, *P* = 0.077; 40 mg/kg: *F*_3,26_ = 0.923, *P* = 0.444) behavior compared to saline control. Bars represent mean ± SEM.

**Supplemental Fig. 5:**
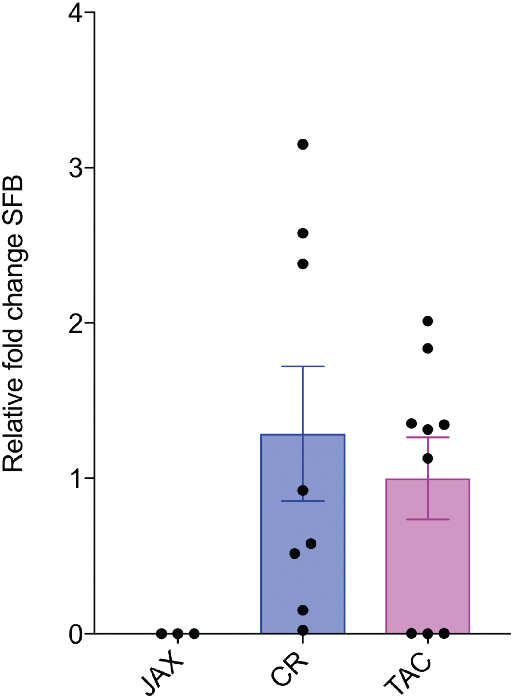
Mice sourced from CR and TAC harbor segmented filamentous bacteria (SFB), while mice sourced from JAX do not. SFB expression was detected using qPCR from the fecal samples from CR and TAC mice. Consistent with previous reports, we did not detect SFB in the fecal samples of mice sourced from JAX. Bars represent mean ± SEM, n > 3 mice per condition.

## Acknowledgments

We wish to thank Dr. David C. Katz for consultation on statistics and editing the manuscript. This work was supported by the Stanley & Jacqueline Schilling Neuroscience Postdoctoral Research Fellowship (M.L.E.), Dennis Weatherstone Predoctoral Fellowships from Autism Speaks (#7825 M.L.E.), the Letty and James Callinan and Cathy and Andrew Moley Fellowship from the ARCS Foundation (M.L.E.), a Dissertation Year Fellowship from the University of California Office of the President (M.L.E.), the Daniel T. O’Connor, MD Memorial Research Grant Scholar Award (J.P.A.), the Eugene Cota- Robles Fellowship (K.P.), P50-MH106438-01 (C.S.C), the University of California Davis Research Investments in Science and Engineering Program (A.K.M.), the MIND Institute IDDRC grant (U54HD079125), and by a grant from the Simons Foundation (SFARI #321998, A.K.M.).

## Financial Disclosures/Conflicts

The authors declare no competing financial interests.

## Supplemental note 1

The microbiome influences disease-relevant outcomes and response to therapeutics (1–3). This source of variability has recently been implicated in susceptibility to MIA outcomes and has relevance for human disease (4–6). Differences in the microbiome may partially account for lack of reproducibility within the MIA model. However, microbiome composition is just one of several common variables rarely tested and known to influence disease-relevant outcomes: these include caging systems (7), shelf level (8), bedding (9), temperature and humidity (10), sex of handler (11), and exposure to pathogens (12). A recent study compared IVC to open cage (OC) systems in the MIA model. Intriguingly, dams housed in IVC, but not OC system, showed a dose-dependent increase in spontaneous abortion (13). This increase in litter loss associated with IVC system was accompanied by increases in maternal and fetal pro-inflammatory cytokine responses to poly(I:C), and changes in the susceptibility to MIA-induced behavioral outcomes. While other husbandry conditions have not been directly evaluated for their effect on MIA outcomes, all the variables listed above alter immune system function. Thus, reporting these variables is paramount for increasing transparency and rigor and improving reproducibility across different research laboratories within the field (14).

## Supplemental Note 2

Lot to lot and even bottle to bottle differences in the composition and preparation of poly(I:C) contribute to variability in the maternal immune response (1, 2). Despite controlling for these factors, we observed a large distribution in the immune response of dams as measured by IL-6 response. Previous reports suggest that the degree of MIA is associated with MIA outcomes (3–5). Thus, individual differences in immune system response to poly(I:C) could lead to heterogeneity in MIA outcomes. To test the baseline immune response (BIR) of our female breeding stock, we injected 7-8 week-old virgin females with a low dose of poly(I:C) and measured serum IL-6 levels 2.5 hr after injection. We used a 4 ± 0.2 mg/kg dosage to assess BIR for all putative dams used to generate offspring for behavioral assays. The variation in dosage is due to the degree of accuracy of the 1 ml insulin syringes used for injections. The cohort was partitioned into BIR groups based upon the cohort distribution, designating the first quartile as ‘low,’ the second and third quartiles as ‘medium,’ and the fourth quartile as ‘high’ responders. Using quartiles to partition our breeding females allowed us to combine results accumulated over multiple lots of ‘priming’ poly(I:C), which differed in the absolute levels of induced IL-6. However, relative demarcations between BIR groups precludes statistical analyses, which require a continuous scale. Thus, we recommend purchasing enough poly(I:C) to complete all experiments within a study so that absolute levels of IL-6 can be compared and analyzed rather than using the BIR grouping strategy employed in this study.

The absolute levels of IL-6 induced by low dose poly(I:C) in virgin females may be influenced by stress (glucocorticoid levels), circadian rhythms, and estrus cycle. While we did not control for these factors in this study, we ensured that the BIR of potential dams was sufficiently stable insofar as the relative categorization (i.e. BIR grouping) did not change when we challenged virgin females a second time (**Supplemental Figure 4**). If comparing absolute levels of IL-6, we recommend controlling for time of injection, estrus cycle, and minimizing stressful vivarium conditions (i.e. cage changing, construction, unfamiliar handlers, etc.), which may influence immune responses.

